# Divergent leaf shapes among *Passiflora* species arise from a shared juvenile morphology

**DOI:** 10.1101/067520

**Authors:** Daniel H. Chitwood, Wagner C. Otoni

## Abstract

**PREMISE OF THE STUDY:** Not only does leaf shape vary between *Passiflora* species, but between sequential nodes of the vine. The profound changes in leaf shape within *Passiflora* vines reflect the temporal development of the shoot apical meristem from which leaves are derived and patterned, a phenomenon known as heteroblasty.

**METHODS:** We perform a morphometric analysis of more than 3,300 leaves from 40 different *Passiflora* species using two different methods: homologous landmarks and Elliptical Fourier Descriptors (EFDs).

**KEY RESULTS:** Changes in leaf shape across the vine are first quantified in allometric terms; that is, changes in the relative area of leaf sub-regions expressed in terms of overall leaf area. Shape is constrained to strict linear relationships as a function of size that vary between species. Statistical analysis of leaf shape, using landmarks and EFDs, reveals that species effects are the strongest, followed by interaction effects, and negligible heteroblasty effects. The ability of different nodes to predictively discriminate species and the variability of landmark and EFD traits at each node is then analyzed. Heteroblastic trajectories, the changes in leaf shape between the first and last measured leaves in a vine, are then compared between species in a multivariate space.

**CONCLUSION:** Leaf shape diversity among *Passiflora* species is expressed in a heteroblastic-dependent manner. Leaf shape is constrained by linear, allometric relationships related to leaf size that vary between species. There is a strong species x heteroblasty interaction effect for leaf shape, suggesting that different leaf shapes between species arise through changes in shape across nodes. The first leaves in the series are not only more like each other, but are also less variable across species. From this similar, shared leaf shape, subsequent leaves in the heteroblastic series follow divergent morphological trajectories. The disparate leaf shapes characteristic of *Passiflora* species arise from a shared, juvenile morphology.

## INTRODUCTION

Heteroblasty is a phenomenon that results from the temporal development of the shoot apical meristem, creating successive changes in the traits of the lateral organs it produces at each node, including the shapes of leaves. Johann Wolfgang von Goethe described the transformation in leaf shape across a shoot as a “metamorphosis”, correctly suggesting that lateral organs are serially homologous structures, from the juvenile and adult leaf shapes a plant displays during vegetative development to reproductive organs (Goethe, 1952; Friedman and Diggle, 2011; Chitwood and Sinha, 2014). There have been many hypotheses about the origins of heteroblastic changes in leaf shape. Inspired by Ernst Haeckel, it was hypothesized that juvenile leaf shapes represented the ancestral condition, and that adult leaves were derived leaf forms (Cushman, 1902; Cushman, 1903). In parallel, another school of thought led by Karl Goebel favored a more environmental explanation for heteroblasty: only supported by the photosynthesis of cotyledons and young leaves, the shape of juvenile leaves represented the aborted development of mature leaf morphology (Goebel, 1908). Careful morphological analysis of young leaf primordia refutes such an idea in Cucurbits (Jones, 1992; Jones, 1995), but classical (Allsopp, 1953a; Allsopp, 1953b; Allsopp, 1954; Njoku, 1956; Njoku, 1971; Roebbelen, 1957; Feldman and Cutter, 1970) and molecular experiments (Yang et al., 2013; Yu et al., 2013) in other species suggest that sugar can serve as a signal hastening the heteroblastic progression of leaf forms.

*Passiflora* species exhibit dramatic heteroblastic changes in leaf shape (**Fig. 1**). *Passiflora edulis* begins with an ellipsoid leaf shape, that transitions to a tri-lobed leaf later in the series (**Fig. 1A**). Like *P. edulis*, *P. caerulea* leaves begin ellipsoid, but transition to a highly-dissected tri-lobe leaf morph and then to four- and five-lobed leaves, and even seven-lobed leaves, as previously reported (Allsopp, 1967) and observed by the authors. Sometimes the transition to lobed leaves is erratic, as in *P. racemosa*, in which the lobes manifest asymmetrically or can revert to the juvenile form with no lobes later in the leaf series (**Fig. 1A**). Other *Passiflora* species exhibit variable degrees of heteroblastic changes in leaf shape (**Fig. 1B**). Unlike the generalized theories of heteroblastic changes in leaf shape inspired by Haeckel or put forth by Goebel, a specific hypothesis has been proposed for the dramatic changes in leaf shape observed in *Passiflora*. Just as diversity in leaf shape among *Passiflora* species is hypothesized to result from diversifying selective pressure from egg-laying *Heliconius* butterflies using leaf shape as a cue (Klucking, 1992; MacDougal, 1994; Gilbert, 1975; Dell’aglio, 2016) diverse leaf shapes in a single vine, resulting from heteroblasty, are thought to similarly deceive butterflies by mimicking non-host plants during critical stages of the vine lifecycle (Gilbert, 1982).

**Figure 1:**
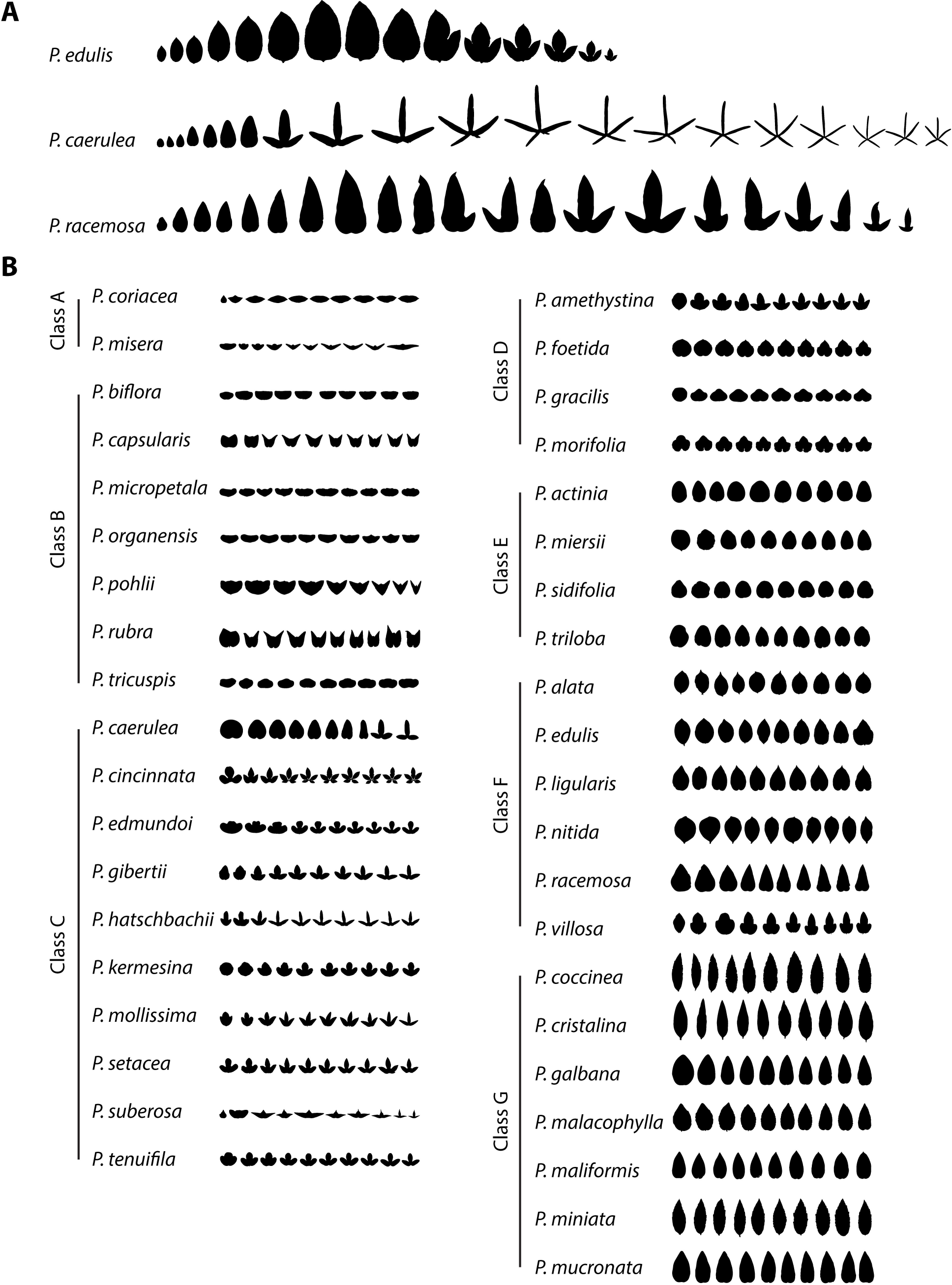
The heteroblastic series in 40 *Passiflora* species. **A)** Notable examples of heteroblastic changes in leaf shape in the indicated *Passiflora* species. **B)** Examples of changes in leaf shape across the heteroblastic series of the 40 *Passiflora* species analyzed in this manuscript, grouped by class. Leaves are scaled such that leaves in the series have the same height. Silhouettes of the first ten leaves of the series are shown. Leaves are arranged from the first leaf (at the shoot base) onwards towards the shoot tip.

Morphometric approaches are critical to separating shape attributes that differentiate species (or genotypes, in genetic studies) regardless of developmental context (Chitwood et al., 2013; Chitwood et al., 2014a) versus purely heteroblastic changes in leaf shape that are shared among closely related species (Chitwood et al., 2012; Chitwood et al., 2016a). Most studies focus on genotype effects that alter leaf shape regardless of node position in the shoot, but the genetic basis of natural variation in heteroblastic shape change itself can also be measured (Chitwood et al., 2014b). The shape features that vary between species are often distinct from those that differentiate leaves arising from sequential nodes, such that using discriminant analyses, species identity can be predicted regardless of node position and vice versa (Chitwood et al., 2016a; Chitwood et al., 2016b). This result is also true among *Passiflora* species (although as later mentioned, it is restricted to certain nodes along the shoot; Chitwood and Otoni, 2016a), demonstrating distinct contributions of species identity and heteroblasty to the shape of each leaf.

Here, we analyze more than 3,300 leaves from 40 *Passiflora* species (Chitwood and Otoni, 2016a; 2016b) but focus on heteroblastic changes in leaf shape. Superimposing averaged leaves from across species, the first two leaves of the series are highly similar in their shape, and more deeply lobed leaves arise later in the the leaf series. Leaf shape changes across the heteroblastic series are allometrically constrained to linear relationships dependent on leaf size, but these allometric constraints vary between species. A statistical analysis of leaf shape shows that species effects are the largest followed by species x heteroblasty interaction effects; heteroblasty effects are negligible. How such strong species x heteroblasty effects manifest among *Passiflora* species is investigated further. Discriminant analysis of shape features capable of predicting species identity for leaves at each node demonstrates that leaves arising from the first nodes at the base of the vine are more like each other and more easily confused between species. The similarity in shape between leaves arising from early nodes between species is supported by an observed lack of variability in juvenile compared to adult leaves. Analysis of changes in shape between the first and last leaves of a vine in a multivariate space suggest that juvenile leaves between species arise from a shared leaf shape. The diversity of leaf shapes observed among *Passiflora* species arises from divergent heteroblastic changes in leaf shape away from a shared, juvenile leaf form.

## MATERIALS AND METHODS

### Data Description

Portions of these Materials and Methods are recapitulated from Chitwood and Otoni (2016a). The two manuscripts analyze different biological problems arising from the same dataset (Chitwood and Otoni, 2016b). Recapitulating the methods here is meant to aid the reader and is in line with Committee on Publication Ethics (COPE) guidelines.

The 555 original scans used for analysis (Chitwood and Otoni, 2016b) represent 40 different species of *Passiflora* in which the order of leaves arising from the vine is recorded (starting with “1” for the youngest leaf scanned from the growing tip of each vine). We importantly note: the numbering of nodes in the raw scans described above (Chitwood and Otoni, 2016b), starting at the tip of the shoot, is opposite from the numbering of nodes presented in the manuscript, in which numbering (starting with “1”) begins with the oldest leaf at the base of the shoot. The reason for this opposite numbering in the manuscript is that by beginning the counting of nodes with “1” at the shoot base the numbering aligns with the heteroblastic series (which begins with the first emerged leaf).

Data from more than 3,300 leaves, and the code to analyze the data and reproduce the figures in this manuscript, are provided, including both landmark data (found in the GitHub repository under PassifloraLeaves/Paper2/Figure2/1.all_landmarks/0.refromated_Proc_coord.txt), measuring the vasculature, lobes, and sinuses, and Elliptical Fourier Descriptor (EFD) data (found in the GitHub repository under PassifloraLeaves/Paper2/Figure2/0.harmonics/0.passiflora_nef.txt), which quantifies leaf outlines (Chitwood, 2016).

### Plant materials and growth conditions

*Passiflora* germplasm was kindly provided by R. Silva (Viveiros Flora Brasil, Araguari, MG, Brazil), Dr. F.G. Faleiro (EMBRAPA Cerrados, Planaltina, DF, Brazil), Prof. M.M. Souza (Universidade Estadual de Santa Cruz - UESC, Ilhéus, BA, Brazil), M. Peixoto (Mogi das Cruzes, SP, Brazil), Prof. M.L. Silva (Universidade do Estado de Mato Grosso, Tangará da Serra, MT, Brazil), and Prof. C.H. Bruckner (Universidade Federal de Viçosa, Viçosa, MG, Brazil).

The plants were germinated from seed, planted between late October 2015 and early March 2016, in Viçosa, at the Federal University of Viçosa, MG, Brazil. The populations were raised and maintained under polycarbonate-covered greenhouse conditions, equipped with automatic environmental control using exhaust fans and evaporative cooling panels (with expanded clay wettable pads). Seeds for each *Passiflora* species were sown in 128 cell propagation plastic trays (GPlan Comércio de Produtos Agrícola s EIRELI – ME, São Paulo, SP, Brazil) filled with horticultural organic Tropstrato HT Hortaliças substrate (Vida Verde Indústria e Comércio de Insumos Orgânicos Ltda, Mogi Mirim, SP, Brazil). After germination (30-40 days), plantlets were individually transplanted to 5 L capacity plastic pots (EME-A-EME Ind. Com. Ltda., Petrópolis, RJ, Brazil) filled with horticultural substrate. Each pot received 5 g of Osmocote^®^ Plus NPK 15-09-12 3-4 month controlled release fertilizer (Scotts, USA). Plants were irrigated on a daily-basis with tap water, and no phytosanitary control was applied.

For scanning, a multifunction printer (Canon PIXMA MX340 Wireless Office All-in-One Printer, model 4204B019, USA) was used. A 20 cm metallic ruler was positioned at the bottom of each scanned sheet as a size marker. Leaves were carefully detached, from the base to the tip of the shoot, and affixed to an A4 paper sheet, adaxial face down, using 12 mm-double sided tape (Scotch Model 9400, 3M do Brasil, SP, Brazil). The numbers written near each leaf indicate position in the shoot, in a tip-to-base direction, starting with the youngest leaf at the tip of the shoot.

### Morphometric and statistical analyses

All morphometric data and code used for statistical analysis is available on GitHub (Chitwood, 2016). Landmarks, as described in the text, were placed on leaves in ImageJ (Abramoff, Magalhaes, and Ram, 2004). Procrustes superimposition was performed using the shapes package (Dryden, 2015) in R (R Development Core Team, 2016) with the procGPA function using reflect=TRUE.

To isolate outlines for Elliptical Fourier Descriptor (EFD) analysis, the “Make Binary” function in ImageJ (Abramoff, Magalhaes, and Ram, 2004) was found to be sufficient to segment leaves. The wand tool was used to select individual binary leaf outlines, which were pasted into a new canvas, which was subsequently saved as an individual image, which was named by vine and node position from which the leaf was derived. The binary images were batch converted into RGB .bmp files and read into SHAPE, which was used to perform chain-code analysis (Iwata et al., 1998; Iwata and Ukai, 2002). The resulting chain-code .chc file was then used to calculate normalized EFDs. The resulting normalized EFD .nef file was then read into Momocs (version 0.2-6) (Bonhomme et al., 2014) in R. The harmonic contributions to shape were visualized using the hcontrib function. Averaged leaf outlines were calculated using the meanShapes function.

Unless otherwise noted, all visualization was performed using ggplot2 in R (Wickham, 2009). Analysis of Variance (ANOVA) was performed using the aov function fitting the model trait ∼ species*heteroblasty. Linear Discriminant Analysis (LDA) was performed using the lda function and subsequent prediction of species identity or heteroblastic node position performed using the predict function with MASS (Venables and Ripley, 2002). For prediction, LDA was performed with CV=”TRUE”, which is a “leave one out” cross-validation approach in which for each leaf an LDA is performed excluding the leaf after which it is assigned to the resulting LDA space it was excluded from. Hierarchical clustering was performed using the hclust function. t-distributed Stochastic Neighbor Embedding (t-SNE) was performed using the Rtsne package (Krijthe, 2015) in R with perplexity=40.

## RESULTS AND DISCUSSION

### Heteroblastic changes in Passiflora leaf shape

The greater than 3,300 leaves analyzed in this study come from 40 different species of *Passiflora* and were collected in order from the base of the vine onwards to the growing tip, keeping track of the node from which each leaf originated. Among the 40 *Passiflora* species sampled, the degree of heteroblastic shape change is variable (**Fig. 1**). Previously (Chitwood and Otoni, 2016a), we defined seven species groups (classes) based on their clustering in a Principal Component Analysis (PCA) and qualitative shape differences. Members of Class G, with lance-shaped leaves, generally exhibit little to no changes in leaf shape across the leaf series, and similarly Class E leaves remain round regardless of shoot position. Members of Class C and D, which are characterized by different numbers of lobes later in the series, begin with a rounder, less lobed juvenile leaf shape at the base of the vine. Members of Class A and B, with wide “bat-like” leaves begin with a rounder leaf shape that transitions to a stereotypical wide leaf (**Fig. 1**).

Generally, if heteroblastic changes in leaf shape are observed, the juvenile leaf form (at the base of the shoot) tends to be rounder and exhibit less lobing compared to adult leaves at the shoot tip (**Fig. 1B**). To substantiate this statement, we used 15 landmarks measuring vascular patterning and the position of the lobes and sinuses, and Elliptical Fourier Descriptors (EFDs), quantifying the contours of leaves, to visualize shape changes through the heteroblastic series (**Fig. 2A-B**). Landmarks are x,y coordinates that define the position of homologous points found on every leaf, such that leaf shape is represented as a set of x,y points shared between samples (Bookstein, 1997). EFDs convert shapes into chain code, a string of numbers that records pixel movements to perfectly recapitulate the shape (Freeman, 1974). The chain code is treated as a wave function and decomposed into a harmonic series using a Fourier transform (Kuhl and Giardina, 1982). The coefficients of the harmonic series represent shape and are used in subsequent analyses.

**Figure 2:**
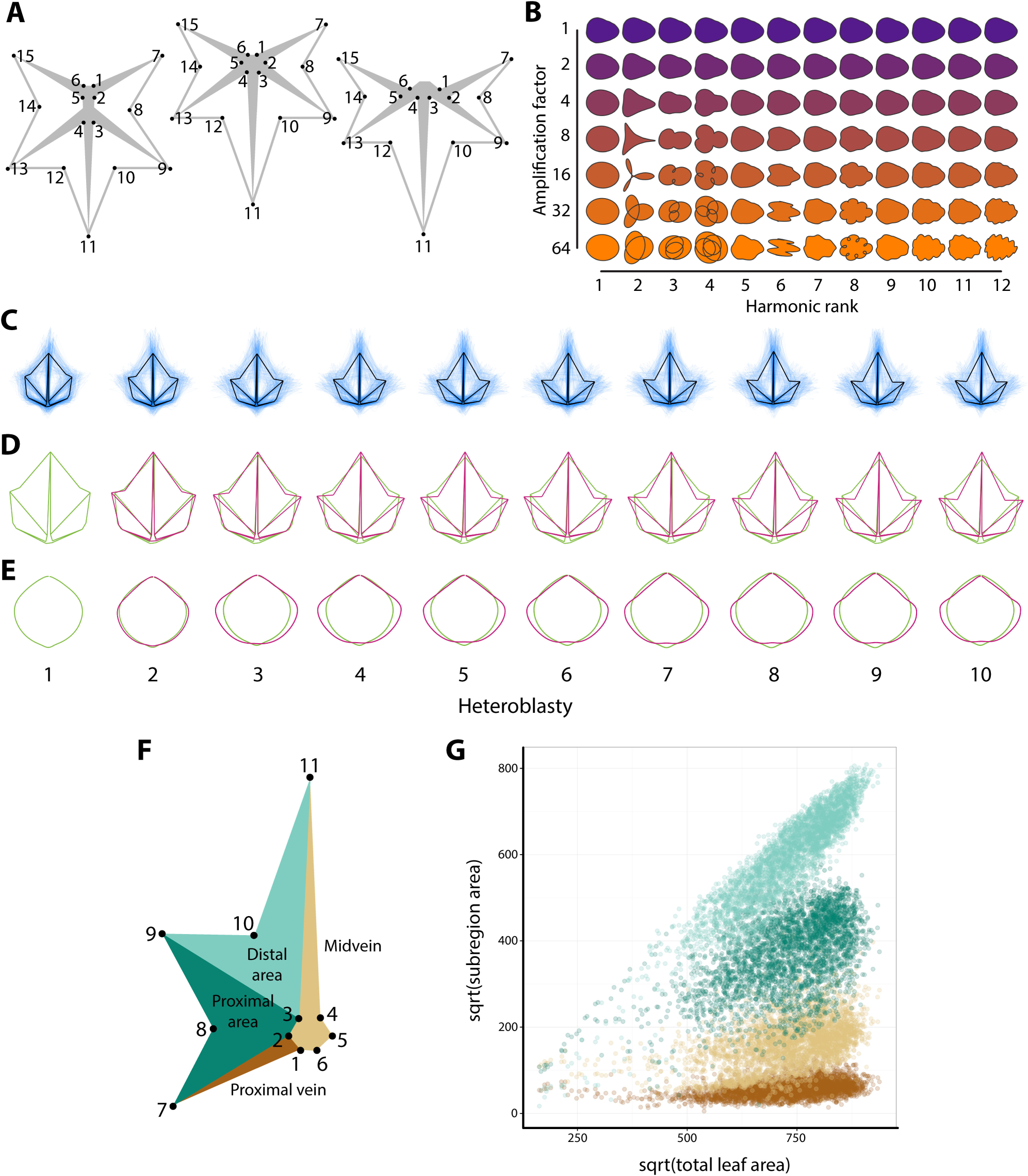
Morphometric methods used to study heteroblastic changes in *Passiflora* leaf shape. **A)** The 15 landmarks used for analysis. Left to right, landmark placement when the distal and proximal veins l) pinnately emerge from the midvein, m) both originate from the petiolar junction, or r) the proximal vein branches from the distal. **B)** Harmonic contributions to shape resulting from Elliptical Fourier Descriptor (EFD) analysis. The harmonic rank is arranged horizontally and the amplification factor vertically. **C)** For each heteroblastic node, the mean leaf as measured with landmarks in shown in black, whereas all landmark data for leaves from the node are depicted in semi-transparent blue. **D)** The average landmark leaf from node 1 is depicted in green and superimposed upon the averaged landmark leaves from other nodes depicted in magenta. **E)** Mean leaves calculated for each heteroblastic node from the harmonic series resulting from an Elliptical Fourier Descriptor (EFD) analysis of leaf contours. The mean contour of leaves from node 1 is depicted in green and the mean contour leaves from other nodes in magenta. **F)** Sub-areas of Procrustes-aligned landmark data calculated for each leaf. **G)** Overall allometric relationships for the square root of distal blade area (light green), proximal blade area (dark green), midvein area (light brown), and proximal vein area (dark brown) plotted against the square root of overall leaf area. All areas are calculated from Procrustes-aligned landmark data as indicated in F). Heteroblastic node position is numbered “1” starting from the shoot base. Note: for convenience to the reader, panels A) and B) are recapitulated in the companion manuscript (Chitwood and Otoni, 2016a). Leaf depicting sub-areas shown in **Fig. 3** re-drawn here for convenience.

Averaged landmarked leaves (**Fig. 2C-D**) and EFD-derived outlines (**Fig. 2E**) across the leaf series compared to the first leaf show that leaves become progressively more dissected. However, most of these heteroblastic changes occur between leaves 1 and 2 at the shoot base, and the remainder of leaves later in the shoot, on average, shows little further changes in shape. This result is consistent with the previous observation (Chitwood and Otoni, 2016a) that shape features in juvenile leaves in the first nodes allow them to be correctly assigned to the predicted node at higher rates than leaves later in the series. We return to this idea later, and explore the hypothesis that across species juvenile leaves are more like each other than leaves later in the series that exhibit divergent shapes.

### Allometric changes in leaf shape across the heteroblastic series

To explore the relative contributions of leaf sub-areas to differences in leaf shape across the heteroblastic leaf series in different species, we performed an allometric analysis. Changes in leaf shape, whether between species or within the heteroblastic series, often correlate linearly with size (Chitwood et al., 2016b); whether such relationships exist in this dataset remains an open question. Procrustes-aligned leaves were divided into sub-regions (**Fig. 2F**), including the areas of the midvein and proximal vein, and distal and proximal leaf blade areas. The rationale of using Procrustes-aligned leaf shapes (which have been scaled, translated, and rotated to superimpose leaves for analysis) is to analyze the relative contributions of vein and blade area to total leaf area. When plotting the square root of each sub-region area against the square root of the overall Procrustes-aligned leaf area, clear linear relationships are observed (**Fig. 2G**). Notably, blade areas expand at a higher rate compared to vein areas as the total leaf area increases (that is, the slope of the blade sub-regions is greater than the vein sub-regions). That total area of the leaf occupied by blade expands at a faster rate than the area occupied by veins is consistent with the previous results observed in grapevines (Chitwood et al., 2016a; Chitwood et al., 2016b).

When the results are plotted for each species, the general trend of blade area expanding at the expense of vein area is observed, but there is a large amount of variation between species (**Fig. 3**). For example, *P. misera* exhibits linear relationships in which both blade regions have similar slopes that are greater than the vein regions. Contrastingly, the distribution of total leaf area for *P. caerulea* is bimodal, and differences in each population of leaves contributes to widely different slopes between the distal and proximal blade sub-regions. The distinctness of each *P. caerulea* sub-population reflects the discrete transformation of leaf shape from entire to highly dissected and palmate (**Fig. 1A**). This is reflected when the heteroblastic node number is projected onto the plots (**Fig. 4**), revealing distinct populations of juvenile and adult leaves with different ratios of blade and vein areas that contribute to each leaf type. Other species vary in the extent that the heteroblastic series is defined by the linear allometric relationships contributing to differences in leaf shape across the leaf series.

**Figure 3:**
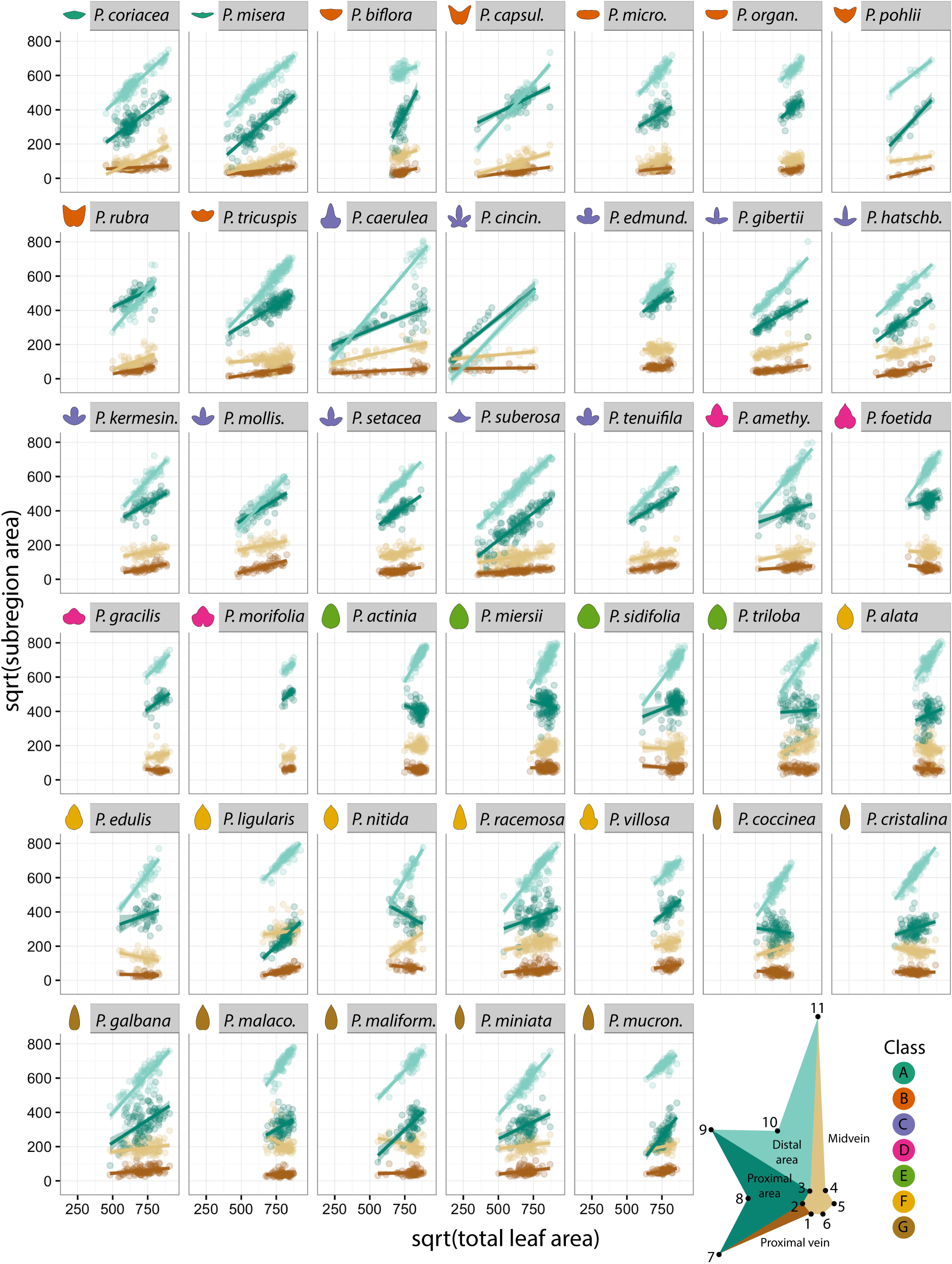
Allometric changes in relative leaf areas in different *Passiflora* species. For each species, the linear relationships between the square root of sub-areas (distal blade area, light green; proximal blade area, dark green; midvein area, light brown; and proximal vein area, dark brown) are plotted against the square root of total area for Procrustes-aligned landmark data. Fitted linear models are super-imposed with 95% confidence bands against data points. Mean leaf contours for each species are provided for reference, colored by class membership. Leaf depicting sub-areas shown in **Fig. 2** redrawn here for convenience.

Although overall blade sub-regions expand at faster rates compared to vein sub-regions, and this relationship is mostly linear across all species (**Fig. 2G**), species vary widely in the relative ratios of these regions across allometric lines (**Fig. 3**) and the heteroblastic series (**Fig. 4**).

**Figure 4:**
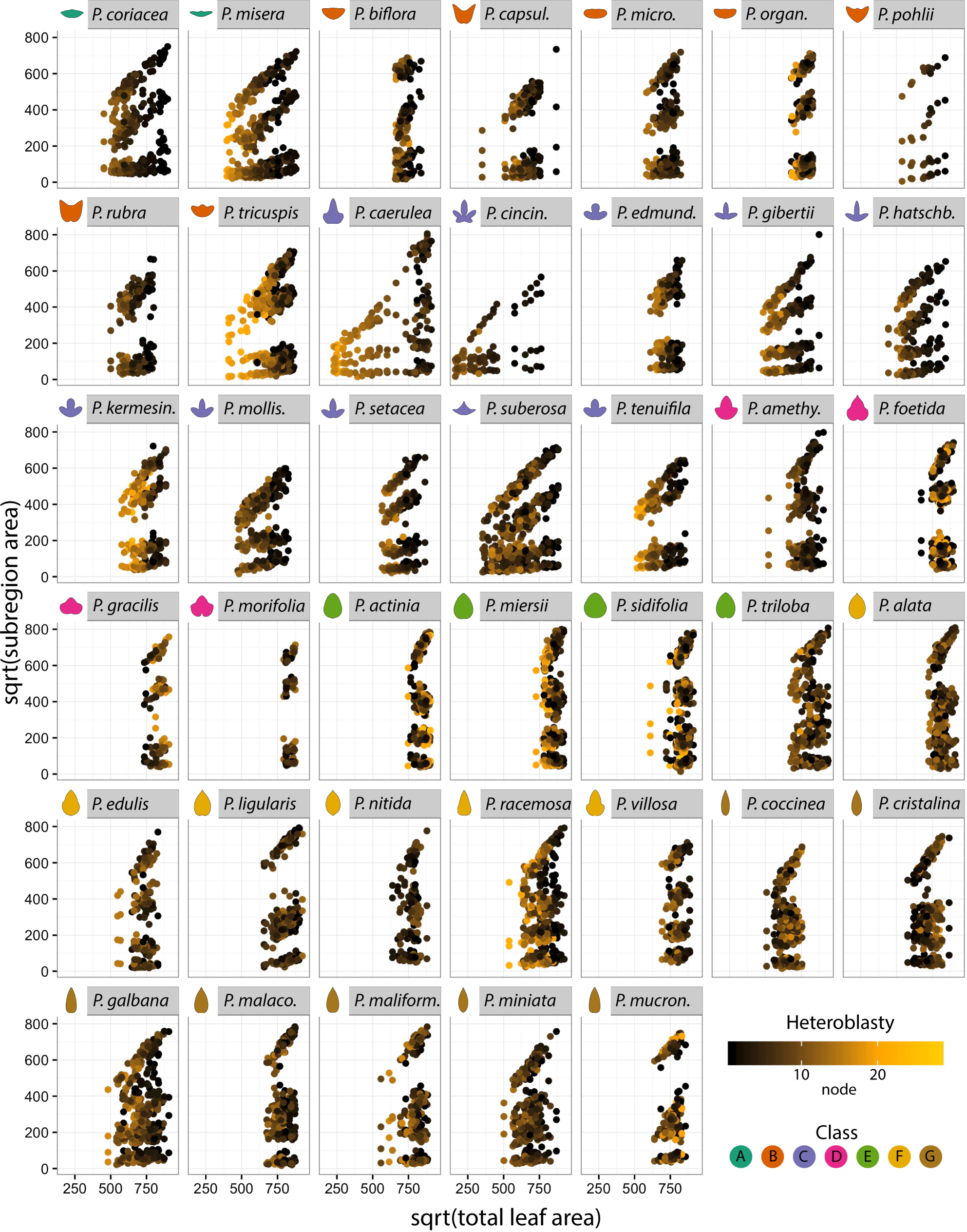
Allometric changes in relative leaf areas across the heteroblastic series. Same plots as in **Fig. 3** except colored by heteroblastic node. For some species, strong linear allometric changes across the heteroblastic leaf series are observed. Heteroblastic node color scheme: shoot base, black; middle shoot, orange; shoot tip, yellow. Heteroblastic node position is numbered “1” starting from the shoot base.

### Statistical effects of species, heteroblasty, and interaction

Previously, we demonstrated using a Linear Discriminant Analysis (LDA) that not only can the leaves of *Passiflora* species be predicted regardless of the node from which they arise, but surprisingly too that the leaf node can be predicted regardless of the species (Chitwood and Otoni, 2016a). This result is consistent with work in grapevine (Chitwood et al., 2016a; 2016b) and demonstrates that within the complex shapes of leaves, some attributes vary independently by species whereas others by node position.

To more rigorously quantify these effects in *Passiflora*, we performed an Analysis of Variance (ANOVA) for each x and y landmark coordinate as well as coefficients of the harmonic series resulting from an Elliptical Fourier Descriptor (EFD) analysis (**Fig. 2A-B**). Each trait was modeled by species*heteroblasty, which takes into consideration the additive species and heteroblastic effects as well as their interaction (**Figs. 5-6**). *P. actinia* was set as the intercept, primarily because its species epithet is alphabetically first, but conveniently, the round shape of *P. actinia* leaves means that since effect sizes are compared against this species (by definition, *P. actinia* effects sizes are 0) the magnitude of leaves of other species deviating from a round shape is being measured.

The largest effect on leaf shape is species (**Fig. 5**). The strongest effect sizes are seen in x landmark coordinates of the petiolar junction in *P. micropetala* and *P. gracilis*, likely owing to the unique intersection of veins in these species. But more general trends are also observable. The x and y landmarks of the distal sinus and harmonic coefficients from the EFD analysis are widely affected in Class C (lobed species) and Classes A and B (wide, wing-shaped species), reflecting the deviation of these leaf shape classes from the round shape of *P. actinia* leaves.

Heteroblasty and interaction effects (**Fig. 6**) were about a magnitude less than species effects (compare legend between **Fig. 5** and **Fig. 6**). The heteroblasty effect is negligible compared to the interaction effect (**Fig. 6**, see top row). This does not mean there are no shape attributes modulated by node position independent from species effects, as we previously showed node position can be predicted independently from species identity (albeit to a much less degree and most strongly for the first two leaves in the heteroblastic series; Chitwood and Otoni, 2016a). Rather, it indicates that heteroblastic-independent effects on leaf shape are slight compared to species and interaction effects. The strongest species x heteroblasty interaction effects are found in the EFD harmonic coefficients of *P. cincinnata*, the leaves of which are highly lobed. Generally, like species effects (**Fig. 5**), the interaction effects are strongest for the lobed (Class C) and winged (Classes A and B) leaf shapes (**Fig. 6**).

**Figure 5:**
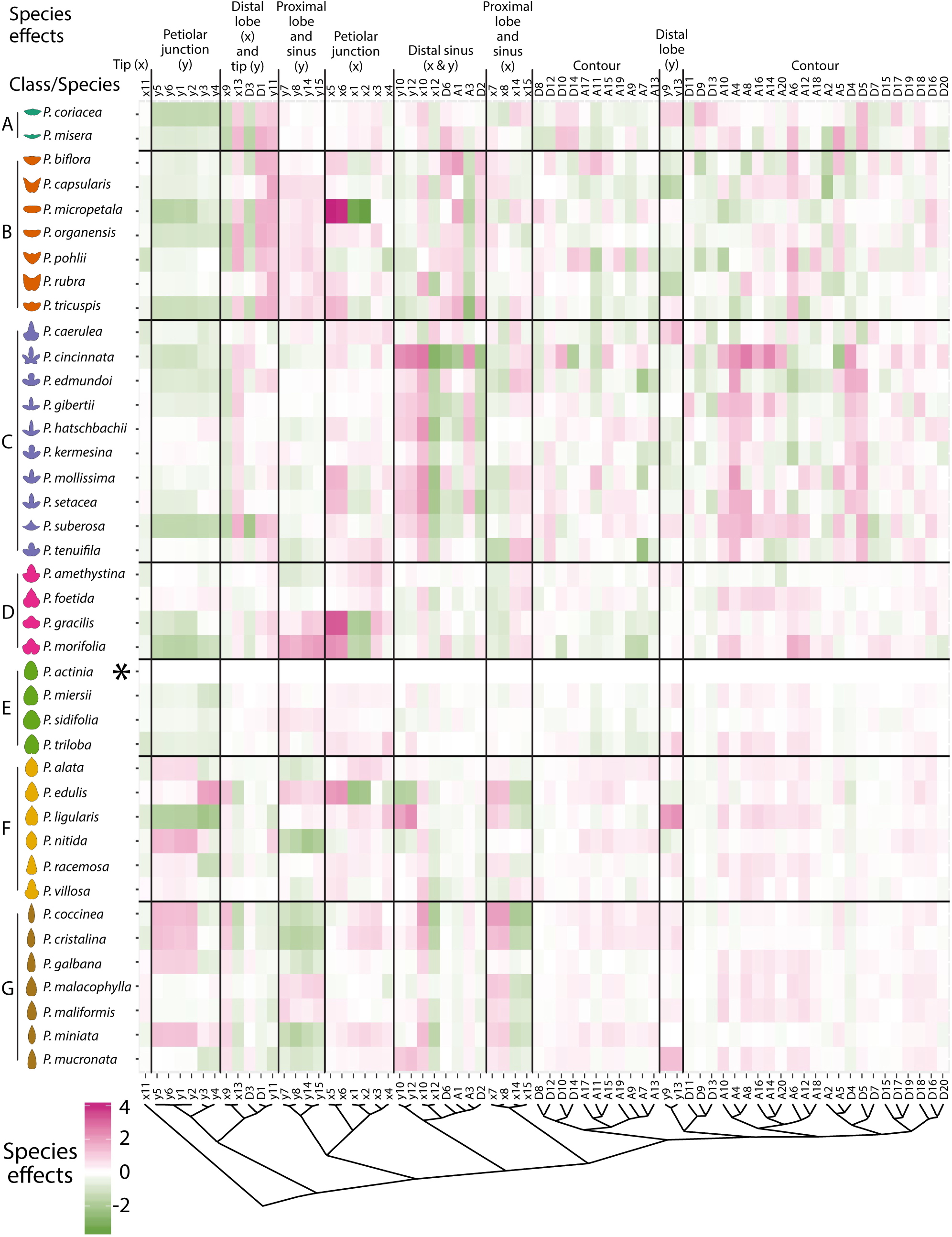
Analysis of Variance (ANOVA) species effects. For each *Passiflora* species separated by class (rows) and x,y landmark and Elliptical Fourier Descriptor (EFD) traits hierarchically clustered (columns) the species effect resulting from the model trait ∼ species*heteroblasty is shown. Opacity indicates magnitude of the species effect and color direction (positive, magenta; negative, green). Effect sizes and directions are relative to *P. actinia* (indicated with asterisk).

**Figure 6:**
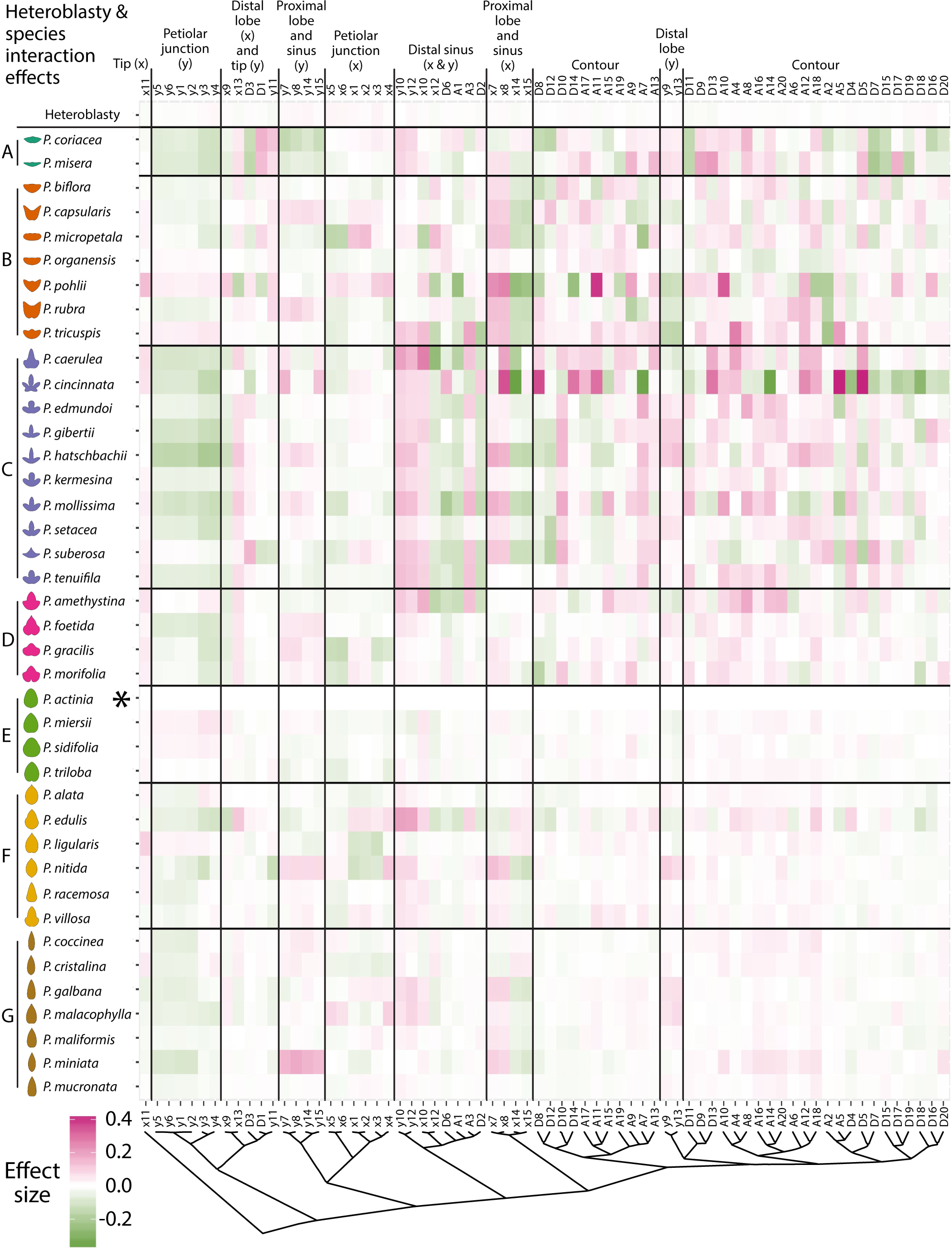
Analysis of Variance (ANOVA) heteroblasty and species x heteroblasty interaction effects. For each *Passiflora* species separated by class (rows) and x,y landmark and Elliptical Fourier Descriptor (EFD) traits hierarchically clustered (columns) the heteroblasty and species x heteroblasty effects resulting from the model trait ∼ species*heteroblasty are shown. The heteroblasty effect is the top row and the species x heteroblasty interaction effects are the subsequent rows. Opacity indicates magnitude of the effect and color direction (positive, magenta; negative, green). Effect sizes and directions are relative to *P. actinia* (indicated with asterisk).

Given that 1) we previously showed that node position can be predicted independently from species identity, although strongest for the first two leaves (Chitwood and Otoni, 2016a) and 2) that the interaction effect sizes between species and heteroblasty are much stronger than heteroblasty alone (**Figs. 5-6**), we hypothesized that evolutionary differences in leaf shape between species of *Passiflora* arise within a heteroblastic context, which we explore more thoroughly, below.

### Divergent heteroblastic trajectories and similar juvenile leaf shapes

Many pieces of evidence suggest the earliest leaf shapes in the heteroblastic series are similar across *Passiflora* species, and that leaves later in the series differ between species. When the average shape of leaves across the heteroblastic series are compared using landmarks (**Fig. 2D**) and contours derived from Elliptical Fourier Descriptors (EFDs) (**Fig. 2E**) it is evident that the more lobed leaf shape characteristic of later leaves is achieved within the first two nodes.

To test the idea that juvenile leaves at the base of the shoot are more similar between species than those later in the series, we performed a Linear Discriminant Analysis (LDA) using both landmark and EFDs to discriminate leaves from each node by species identity (**Fig. 7A**). Although there is wide variability of the ability to discriminate the leaves of each species, the LDAs using leaves from nodes 1 and 2 performed poorly in their ability to discriminate leaves by species compared to subsequent nodes. From the overall average correct reassignment rate for the LDA performed for each node (see bottom of **Fig. 7A**) it is evident that leaves from nodes 1 and 2 have less distinctive features differentiating leaves from species compared to later nodes. This suggests that leaves from nodes 1 and 2 are more similar in shape between species than leaves from later nodes.

**Figure 7:**
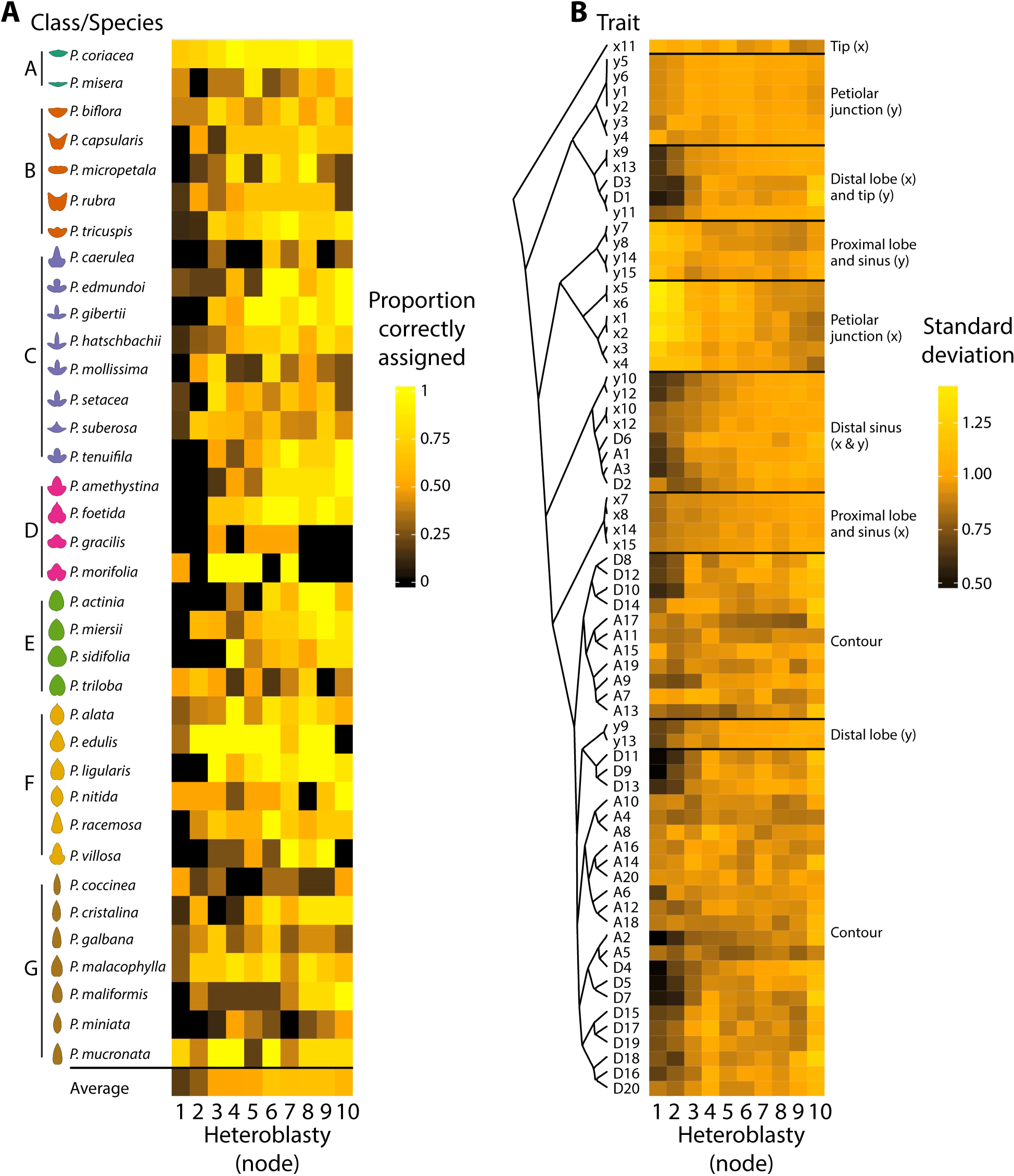
Juvenile leaves are similar in shape across *Passiflora* species. **A)** Heatmap showing the proportion of correctly assigned species from Linear Discriminant Analyses (LDAs) performed with both landmark and Elliptical Fourier Descriptor (EFD) data for each heteroblastic node. The average correct assignment across species is provided as well. Generally, the proportion of leaves correctly assigned to species increased with heteroblastic node number. Averaged contours of leaves from each species are provided for reference and colored by class. **B)** For each trait, the standard deviation across each heteroblastic node is shown. Traits are arranged by hierarchical clustering and groups corresponding to x and y coordinates of landmarks indicated. Proportion correctly assigned: low, black; middle, orange; high, yellow. Standard deviation: low, black; middle, orange; high, yellow. Class color scheme: class A, teal; class B, orange; class C, lavender; class D, magenta; class E, green; class F, yellow; class G, brown. Heteroblastic node position is numbered “1” starting from the shoot base.

A more direct test of variability in leaf shape is to measure the standard deviation of the raw traits used to measure leaf shape. Shape features of the leaf were hierarchically clustered and the standard deviation for each across species for each node calculated (**Fig. 7B**). Most shape features either have reduced or unchanged standard deviation values in the first 1-3 leaves of the series compared to later leaves. A minority of shape features, especially the x coordinate values for landmarks defining the petiolar junction and bases of the major veins (landmarks 1-6) have increased standard deviation values in the earlier nodes. The results suggest that generally leaf shape is less variable in leaves from the first nodes (**Fig. 7B**) consistent with the observation that juvenile leaves discriminate species less than leaves later in the series (**Fig. 7A**). Exceptionally, the landmarks defining the petiolar junction in the x coordinate direction are more variable in juvenile than adult leaves (**Fig. 7B**).

To visualize the divergent heteroblastic trajectories leading to disparate leaf shapes between *Passiflora* species, we used a t-distributed Stochastic Neighbor Embedding (t-SNE) approach to reduce the dimensionality of the data (**Fig. 8A**) (Krijthe, 2015). t-SNE separates species classes and benefits from no assumptions of linearity and reducing the data to strictly two dimensions. By doing so, each sampled vine can be visualized as a vector in two-dimensional space, with the base and tip of the vector corresponding to the first sampled node at the base of the shoot and the furthest sampled node at the tip of the shoot, respectively (**Fig. 8B**). Each vector, therefore, is a representation of the shape space traversed over nodes across the heteroblastic series.

**Figure 8:**
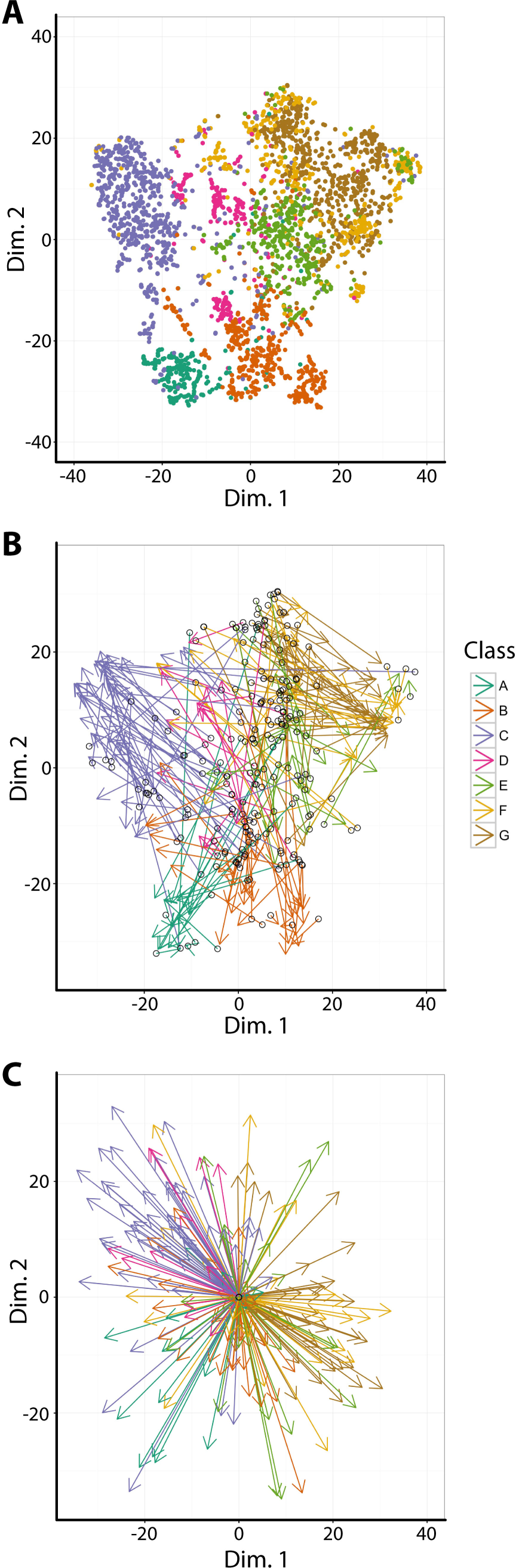
Traversal of the heteroblastic series through t-SNE space. **A)** Graph of Dimension 2 vs. Dimension 1 of leaves, colored by class, in a t-Distributed Stochastic Neighbor Embedding (t-SNE) analysis. **B)** Same data as in A) except with arrows corresponding to the leaf series collected for each plant. The first node (towards the base of the shoot) is the arrow base and indicated with a hollow, black circle. The last node (towards the tip of the shoot) is the arrow head. Arrows are colored by species class. **C)** The same arrow data as in B) except that the base of all arrows have been translated to the origin of Dimension 2 vs. Dimension 1. Arrows are colored by species class. Class color scheme: class A, teal; class B, orange; class C, lavender; class D, magenta; class E, green; class F, yellow; class G, brown.

The bases of the vectors representing each vine tend to cluster together, with similar Dimension 1 values but varying across Dimension 2. From this common region representing a shared juvenile leaf shape, the directions of each vector for different species classes vary, representing differing heteroblastic trajectories leading to disparate adult leaf shapes at nodes towards the shoot tips. To better visualize the divergent heteroblastic trajectories of each species class, each vector base was centered to the origin (**Fig. 8C**). After centering, it is apparent that different species classes vary drastically—sometimes diametrically opposed—in the direction of their heteroblastic shape changes.

Collectively, the inability of leaves from the first nodes to successfully discriminate different species (**Fig. 7A**), the reduced variability in shape features of leaves from the first nodes (**Fig. 7B**), and the divergent heteroblastic trajectories between species classes (**Fig. 8**) demonstrate that juvenile leaves between *Passiflora* species are similar and that divergent heteroblastic trajectories are responsible for the distinctive leaf shapes between species.

## CONCLUSIONS

Many explanations for heteroblasty, the changes in leaf shape and other traits across a shoot resulting from the temporal development of the shoot apical meristem from which lateral organs arise, have been proposed. The idea that juvenile and adult leaves recapitulate the ancestral and derived leaf forms across evolution (Cushman, 1902; Cushman, 1903) or that juvenile leaves result from the lack of photosynthate to complete development (Goebel, 1908) have been proposed as possible explanations for the dramatic changes in leaf shape across a shoot. In *Passiflora*, the heteroblastic series has been hypothesized to be a mechanism to avoid *Heliconius* butterflies that use leaf shape as a cue to lay eggs (Gilbert, 1982). Although we cannot distinguish between these alternatives, it is nonetheless important to quantify changes in leaf shape across the heteroblastic series, so that a rigorous understanding of how leaf shape changes manifest across vines contribute to diversity within the genus *Passiflora* (**Fig. 1**). Doing so provides insight into how different leaf shapes arise within a developmental and evolutionary context.

Superimposing averaged leaf shapes from across the heteroblastic series, the leaves arising from the first two nodes share a similar shape and more deeply lobed leaves arise later in the series (**Fig. 2C-E**). These heteroblastic changes in leaf shape are allometrically constrained to linear relationships, and these strict linear relationships vary between species (**Figs. 2-4**). Statistically modeling species, heteroblasty, and interaction effects, species effect sizes are the largest (**Fig. 5**) followed by interaction effects approximately a magnitude less and negligible heteroblasty effects (**Fig. 6**). Considering the unique similarity in the shape of leaves arising from the first two nodes, we hypothesized that divergent leaf shapes between *Passiflora* species arising later in the series arise from a shared, juvenile leaf shape. Juvenile leaves are more often mistakenly identified between species than adult leaves found later in the shoot (**Fig. 7A**), and consistent with juvenile leaves resembling each other, the variability of most morphometric features is lower in juvenile compared to adult leaves (**Fig. 7B**). Comparing the first and last leaves of a shoot within a multivariate space, the heteroblastic trajectories of different species are divergent, originating from a similar juvenile form but traversing towards disparate shapes (**Fig. 8**). Our data show that the striking differences in leaf shape between *Passiflora* species are expressed in a developmental manner, later in the heteroblastic series, arising from a shared juvenile leaf shape.

## Authors’ contributions

The overall project was conceived by DHC and WCO. WCO grew and scanned all plant material and DHC carried out analysis. DHC and WCO wrote the paper.

